# The cell cycle regulator GpsB functions as cytosolic adaptor for multiple cell wall enzymes

**DOI:** 10.1101/427823

**Authors:** Robert M. Cleverley, Zoe J. Rutter, Jeanine Rismondo, Federico Corona, Ho-Ching Tiffany Tsui, Fuad A. Alatawi, Richard A. Daniel, Sven Halbedel, Orietta Massidda, Malcolm E. Winkler, Richard J. Lewis

## Abstract

Bacterial growth and cell division requires precise spatiotemporal regulation of the synthesis and remodelling of the peptidoglycan layer that surrounds the cytoplasmic membrane. GpsB is a cytosolic protein that affects cell wall synthesis by binding to the cytoplasmic mini-domains of peptidoglycan synthases to ensure their correct subcellular localisation. Here we have discovered critical structural features for the interaction of GpsB with peptidoglycan synthases from three different bacteria and demonstrated their importance for cell wall growth and viability. We have used these structural motifs to predict and confirm novel partners of GpsB in *Bacillus subtilis*, illuminating the role of this key regulator of peptidoglycan synthesis. GpsB thus functions as an adaptor, to mediate the interaction between membrane proteins, scaffolding proteins, signalling proteins and enzymes to generate larger protein complexes at specific sites in a bacterial cell cycle-dependent manner. Given the importance of GpsB in pathogenic bacteria, this study has not only revealed mechanistic details of how cell wall synthesis is co-ordinated with the bacterial cell cycle but could also represent a starting point for the design of much needed new antibiotics.

---

Peptidoglycan (PG), a network of glycan strands connected by short peptides, forms the essential cell envelope that maintains cell shape and protects bacteria from osmotic stresses^1^. High molecular weight (HMW) bi-functional penicillin binding proteins (class A PBPs) are PG synthases that catalyse glycan strand polymerisation and peptide crosslinking, whereas HMW class B mono-functional PBPs only have transpeptidase functions^2^. The PG layer needs remodelling to enable normal cell growth and division and thus the bacterial cell cycle requires the extracellular activities of PBPs^3^ and PG hydrolases^4^ to be co-ordinated. The outer membrane-anchored LpoA/B lipoproteins activate their cognate PBP1A/1B PG synthases in the synthesis of the thin, periplasmic PG layer in the Gram-negative paradigm *Escherichia coli*^5,6^. By contrast, Gram-positive bacteria have a much thicker PG layer that is complemented with other anionic cell wall polymers. PG synthesis regulation in Gram-positive bacteria involves protein phosphorylation by orthologues of the serine/threonine kinase PknB^7^/StkP^8^, and dedicated cell cycle scaffolding proteins including DivIVA^9^, EzrA^10^ and GpsB^11,12^. However, the molecular mechanisms that modulate PG synthesis in Gram-positive bacteria are virtually unknown.

GpsB has emerged as a major regulator of PG biosynthesis in low G+C Gram-positive bacteria, and its homologues (DivIVA/Wag31/antigen 84) in Actinobacteria play essential roles in hyphal growth and branching^13–15^. It was initially characterised in *Bacillus subtilis* where severe cell division and growth defects were observed when both *gpsB* and *ezrA*^11^ or *gpsB* and *ftsA*^12^ were deleted. Both EzrA and FtsA play roles in the dynamics and membrane anchoring of the FtsZ Z-ring, the constriction of which is fundamental to cell division^16^. The Z-ring also recruits downstream proteins, including PBPs^17,8^, to complete the process. Deletion of *gpsB* alone in *Listeria monocytogenes* causes marked growth and division defects at 37°C and is lethal at 42°C^19^. Moreover, *gpsB* deletion in *L. monocytogenes* also results in enhanced susceptibility to β-lactam^19^ and fosfomycin^20^ antibiotics, reduced virulence in an insect infection model^19^, and caused alterations to PG structure^21^. Mutations in *gpsB* that affect binding to the PG synthase PBPA1 also show a lethal phenotype in *L. monocytogenes* at 42°C^19^. The *gpsB* gene is essential in the *Streptococcus pneumoniae* D39 progenitor strain as well as in some of its laboratory derivatives and its inactivation results in elongated cells unable to divide^22–24^. In addition, a recent genome-wide association study of *S. pneumoniae* clinical isolates revealed that the presence of *gpsB* variants is correlated significantly to β-lactam resistance^25^, suggesting that GpsB may have fitness and pleiotropic roles in maintaining cell wall integrity during antibiotic stress.

In both *B. subtilis*^11^ and *L. monocytogenes*^19^ the cytosolic GpsB localises to the lateral side walls of newborn, growing cells and to the septum of dividing cells, the same localisation pattern as that of *B. subtilis* PBP1^11^. In *S. pneumoniae*, GpsB localises to mid-cell^22^, the only region of active PG synthesis for both peripheral (side-wall) elongation and cell division in this bacterium. The localisation of GpsB at regions of active PG synthesis allows for the interaction of GpsB with the poorly characterised cytoplasmic mini-domains of PG synthases^11,19,26,27^. *S. pneumoniae* GpsB (*Sp*GpsB) has been found to co-immunoprecipitate with *Sp*PBP2a, *Sp*PBP2b and *Sp*MreC^24^, suggesting these proteins form a complex that is regulated by *Sp*GpsB^24^.

To gain molecular understanding of GpsB function, we have solved three crystal structures of PBP cytoplasmic mini-domains in complex with GpsB, the first structures of a PG synthase in complex with a cytoplasmic cell cycle regulator. Despite a marked absence of sequence and structural homology, we have discovered that the PBP domains interact with equivalent surfaces in GpsB using an arginine that is conserved in the respective orthologues of the PBPs. The visualisation of each complex has allowed a comprehensive mutagenesis strategy and functional study to rationalise the role of each interfacial amino acid in the PBP:GpsB pairs. We have discovered a sequence motif used by the *B. subtilis* PG synthase to interact with GpsB. This motif has been used to query the *B. subtilis* proteome for potential new partners of GpsB, and we have identified two new members of the GpsB interactome in this organism, and provide evidence for their connection to other, established proteins in growth and division. Therefore, the role of GpsB in the bacterial cell cycle is as an adaptor^28–30^, docking PG synthases to other cell wall enzymes, scaffolds and shape determinants into protein complexes that drive division (divisome) and peripheral growth (elongasome).

## Results

### The first 16 amino acids of *Bs*PBP1 drive the interaction with *Bs*GpsB

GpsB is an influential cell cycle regulator in low G+C Gram-positive bacteria and hence we set out to establish the common rules by which GpsB interacts with major PG synthases in three important bacteria - one model species (*B. subtilis*) and two pathogens (*L. monocytogenes* and *S. pneumoniae*). It had been determined previously by us that full-length *B. subtilis* PBP1 (*Bs*PBP1) bound to full-length *Bs*GpsB with a *K*_d_ of 0.7 μM^19^. Deletion of the first 16 amino acids of *Bs*PBP1_17–914_ severely affected *Bs*GpsB binding: no interaction was observed even when 25 μM *Bs*GpsB was injected over a *Bs*PBP1_17–914_-immobilised SPR chip (**Figure 1A**). We subsequently solved the crystal structure of the *Bs*GpsB_5–64_:*Bs*PBP1_1–17_ complex (**Figure 1B,1C**). The *Bs*PBP1_1–17_ peptide is predominantly α-helical and there are no substantial conformational changes in unbound *Bs*GpsB_5–64_ on binding *Bs*PBP1_1–17_. The *Bs*PBP1_1–17_ α-helix is stabilized by an intramolecular salt bridge between Glu9 and Arg12 and by a hydrogen bond between the sidechain of Ser7 and the backbone amide of Ala10. A prominent feature of the complex is the deep penetration of the sidechain of *Bs*PBP1^Arg8^ into the groove between *Bs*GpsB_5–64_ α-helices 1 and 2, contacting the mainchain carbonyl oxygens of *Bs*GpsB^Ile13^, *Bs*GpsB^Leu14^ and *Bs*GpsB^Lys16^ and forming a salt bridge with *Bs*GpsB^Asp31^ (**Figure 1C**), which in turn is tethered in place by hydrogen bonds to the hydroxyl of *Bs*GpsB^Tyr25^. The backbone amides of *Bs*PBP1^Arg8^ and *Bs*PBP1^Glu9^ interact with *Bs*GpsB^Asp35^ mimicking the mainchain interactions in successive turns in an α-helix. In a longer α-helix, the backbone amides of *Bs*PBP1^Arg8^ and *Bs*PBP1^Glu9^ would not be available to interact with *Bs*GpsB^Asp35^ because of intra-helical hydrogen bonds with the mainchain carbonyls of *Bs*PBP1^Phe5^ and *Bs*PBP1^Asn6^. The sidechain of *Bs*PBP1^Arg11^ forms hydrogen bonds with the carbonyl oxygen of *Bs*GpsB^Leu14^ and a salt bridge with *Bs*GpsB^Glu17^. Van der Waals’ interactions connect *Bs*PBP1^Arg8^ and *Bs*GpsB^Leu34^ (**Figure 1C**), and *Bs*PBP1^Glu9^ and *Bs*GpsB^Lys32^.

**Figure 1:**
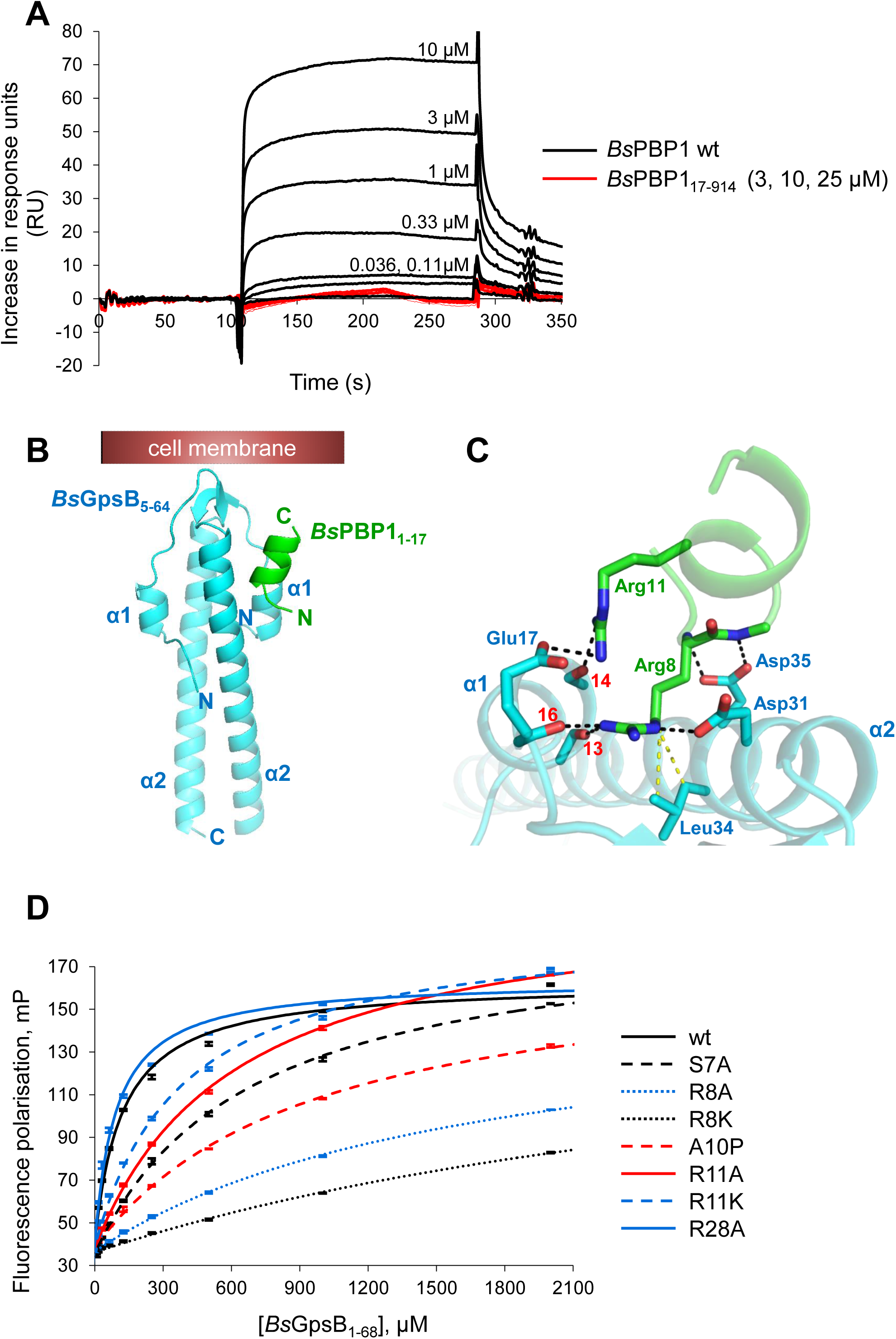
*Bs*GpsB:*Bs*PBP1 interactions are driven by conserved arginines in the α-helical cytoplasmic minidomain of *Bs*PBP1. (**A**) *Bs*GpsB interacts with the first 16 amino acids of *Bs*PBP1. SPR sensorgrams of full-length *Bs*GpsB against immobilised full-length *Bs*PBP1 (black) and *Bs*PBP1_17–914_ (red). *Bs*GpsB does not interact with the *Bs*PBP1_17–914_ coated chip, even when 25 μM GpsB is injected. (**B**) Cartoon of the crystal structure of the *Bs*GpsB_5–64_:*Bs*PBP1_1–17_ complex. *Bs*GpsB_5–64_ is coloured cyan and *Bs*PBP1_1–17_ is coloured green. The *Bs*PBP1_1–17_ peptide binds to a groove between α-helices 1 and 2 in only one molecule of *Bs*GpsB_5–64_ in the crystallographic asymmetric unit as the second *Bs*GpsB-binding site is blocked by crystal contacts. The likely plane of the bacterial membrane is shown as a red box. (**C**) The *Bs*GpsB_5–64_:*Bs*PBP1_1–17_ complex is driven by a conserved SRxxR(R/K) motif in *Bs*PBP1. Key interfacial residues in the *Bs*GpsB_5–64_:*Bs*PBP1_1–17_ complex are shown as sticks and coloured (and labelled) blue and green, respectively. The carbonyl oxygens of *Bs*GpsB^Ile13^, *Bs*GpsB^Leu14^ and *Bs*GpsB^Lys16^ are labelled with their respective red numerals. Hydrogen bonds are shown as black dashed lines and the van der Waals’ interactions between *Bs*GpsB^Leu34^ and *Bs*PBP1^Arg8^ are in yellow. (**D**) Mutation of key *Bs*PBP1 interfacial residues in the structure of *Bs*GpsB_5–64_:*Bs*PBP1_1–17_ leads to a loss of binding of TAMRA-labelled *Bs*PBP1_1–32_ variants to *Bs*GpsB_1–68_ as measured by fluorescence polarisation. The calculated dissociation constants can be found in **Supplementary Table 1**.

The importance of the interactions described above was confirmed by fluorescence polarisation (FP) and circular dichroism (CD). The *Bs*GpsB_1–68_^Glu17Ala^, *Bs*GpsB^Tyr25Phe^, *Bs*GpsB_1–68_^Asp31Ala^ and *Bs*GpsB_1–68_^Asp35Ala^ mutations had little impact on protein stability (**Supplementary Figure 1A**) and each reduced the affinity for *Bs*PBP1_1–32_ by more than 8-fold (**Supplementary Figure 1B, Supplementary Table 1**). *Bs*PBP1^Arg8Lys^, *Bs*PBP1^Arg8Ala^ and *Bs*PBP1^Arg11Ala^ mutations each resulted in reduced affinities for *Bs*GpsB_1–68_ by at least 5-fold (**Figure 1D, Supplementary Table 1**). *Bs*PBP1_1–32_^Arg28Ala^ had no effect on binding (**Figure 1D, Supplementary Table 1**), confirming that non-specific electrostatics do not drive the *Bs*GpsB:*Bs*PBP1 interaction. The Ser7Ala and Ala10Pro mutations each reduced the affinity for *Bs*GpsB_1–68_ by at least 6-fold (**Figure 1D, Supplementary Figure 1C, Supplementary Table 1**) by affecting the α-helix of *Bs*PBP1_1–32_. Ser7 acts as the helix N-cap, a role that can also be performed by Asn and Thr^31^, and substitutions equivalent in helical positions to Ser7Ala and Ala10Pro destabilise model peptides^31,32^. Finally, the importance of PBP1 Ser7, Arg8 and Arg11 to GpsB binding is highlighted because these are the most well conserved residues in an alignment of the cytoplasmic mini-domains of *Bacillaceae* PBP1 PG synthases (**Supplementary Figure 1D**).

### *L. monocytogenes* GpsB interacts with PBPA1 via a conserved TRxxYR motif

The deletion of *gpsB* alone in *B. subtilis* has no readily-apparent phenotype until combined with deletions in *ezrA*^12^ or *ftsA*^12^; by contrast, the deletion of *gpsB* in *L. monocytogenes* is lethal when grown at 42°C^19^. Since GpsB in both species interact with class A PG synthases, we next determined whether the rules established above for the *Bs*GpsB:*Bs*PBP1 interaction could be applied directly to *Lm*GpsB:*Lm*PBPA1. The cytoplasmic mini-domain of *Lm*PBPA1 has an abundance of positively charged residues (**Supplementary Figure 2A**), but lacks an exact copy of the SRxxR(R/K) motif of *Bacillaceae* PBP1 (**Supplementary Figure 1A**), the closest equivalent is TRxxYR. In FP *Lm*PBPA1_1–20_ bound to *Lm*GpsB_1–73_ with an affinity similar to that of *Bs*PBPA1_1–32_ for *Bs*GpsB_1–68_ (**Supplementary Figure 2B, Supplementary Table 1**), but we were unable to co-crystallize *Lm*GpsB with *Lm*PBPA1 peptides to visualise these interactions and to compare them to *Bs*GpsB_5–64_:*Bs*PBP1_1–17_. Consequently we co-crystallized *Bs*GpsB_5–64_^Lys32Glu^:*Lm*PBPA1_1–21_ that can act as a surrogate because (i) all the GpsB interfacial residues in the *Bs*GpsB_5–64_:*Bs*PBP1_1–17_ complex are conserved in *Lm*GpsB except for Lys32, which is glutamate in *Lm*GpsB; (ii) *Lm*GpsB and *Bs*GpsB use overlapping PBP binding sites^19^; (iii) the *K*_d_s of *Bs*GpsB_1–68_^Lys32Glu^:*Lm*PBPA1_1–20_ and *Lm*GpsB_1–73_:*Lm*PBPA1_1–20_ are almost identical (**Supplementary Figure 2B, Supplementary Table 1**). However, the majority of the *Lm*PBPA1_1–15_ peptide in the subsequent structure of the *Bs*GpsB_1–68_^Lys32Glu^:*Lm*PBPA1_1–15_ complex was disordered except for an arginine occupying the same orientation as *Bs*PBP1_1–17_^Arg8^ in the *Bs*GpsB_5–64_:*Bs*PBP1_1–17_ complex (**Figure 2A**). This sole arginine makes the same interactions as described above (**Figure 1C**). The interaction of *Bs*GpsB_5–64_^Lys32Glu^ with *Lm*PBPA1_1–15_ thus centres almost entirely on a single arginine and how *Lm*GpsB discerns *Lm*PBPA1 over other arginine-rich peptides was determined by FP. *Lm*PBPA1_1–20_^Arg8Ala^ and *Lm*PBPA1_1–20_^Arg12Ala^ reduced the affinity for *Bs*GpsB_1–68_^Lys32Glu^ by >15- and ~4-fold, respectively (**Supplementary Figure 2B, Supplementary Table 1**). Reintroducing positive charge into *Lm*PBPA1_1–20_^Arg8Ala^ did not restore wild-type binding affinity as *Lm*PBPA1_1–20_^Arg8AlaSer16Arg^ bound to *Bs*GpsB_1–68_^Lys32Glu^ with an affinity at least 10-fold weaker than wild-type (**Supplementary Figure 2B, Supplementary Table 1**). Bacterial two-hybrid assays (BACTH) support the central importance of *Lm*PBPA1^Thr7^, *Lm*PBPA1^Arg8^ and *Lm*PBPA1^Arg12^, and to a lesser extent *Lm*PBPA1^Tyr11^, and that non-specific electrostatic interactions play little part in binding *Lm*GpsB (**Figure 2B**).

**Figure 2:**
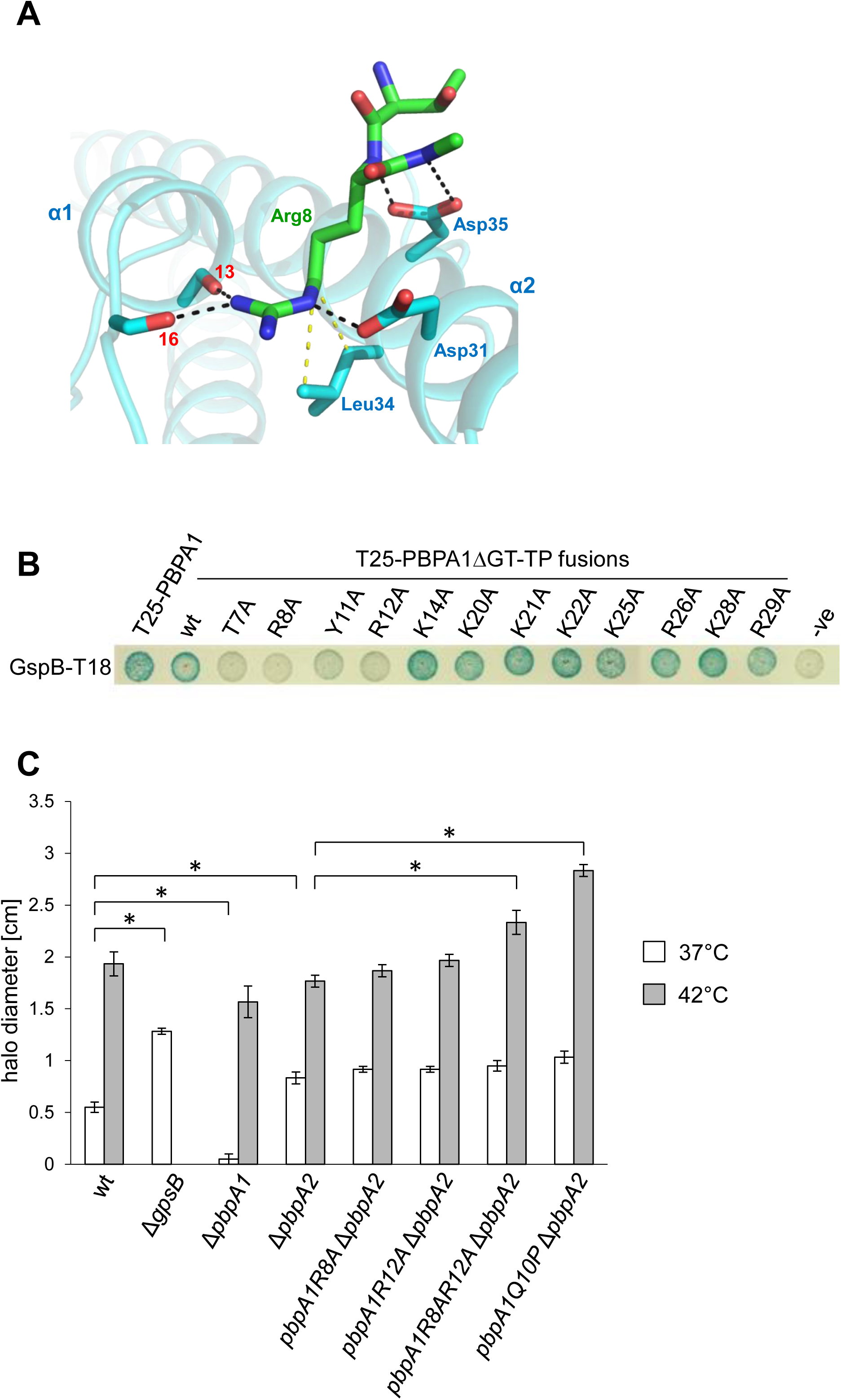
The *Lm*GpsB:*Lm*PBPA1 interactions are also driven by a conserved arginine. (**A**) The structure of the *Bs*GpsB_5–64_^Lys32Glu^:*Lm*PBPA1_1–15_ complex reveals that only Arg8 of the *Lm*PBPA1_1–15_ peptide is ordered. In this cartoon, *Bs*GpsB_5–64_^Lys32Glu^ is coloured cyan and selected sidechains are drawn as stick with cyan carbons, whereas the *Lm*PBPA1_1–15_ peptide is represented in stick form, with green carbons. The carbonyl oxygens of *Bs*GpsB^Ile13^ and *Bs*GpsB^Lys16^ are denoted by respective red numerals. Hydrogen bonds are shown as black dashed lines and the van der Waals’ interactions between *Bs*GpsB^Leu34^ and *Lm*PBPA1^Arg8^ are in yellow. Only one PBP-binding site is occupied by peptide in these crystals because the second site is blocked by crystal contacts. (**B**) Mutation of conserved *Lm*PBPA1 residues results in a loss of interaction by BACTH. The removal of residues 92 to 827, correlating to the glycosyltransferase and transpeptidase domains of *Lm*PBPA1, results in the PBPA1ΔGT-TP peptide. Empty pKT25 (-) was used as a negative control. Agar plates were photographed after 48 hours at 30°C. (**C**) Effect of N-terminal *pbpA1* mutations on fosfomycin sensitivity of a Δ*pbpA2* mutant. Fosfomycin inhibits the first enzyme in the biosynthetic pathway of PG, MurA, and the Δ*gpsB* mutant is hypersensitive to fosfomycin probably because of unproductive consumption of PG precursors due to mis-regulated *Lm*PBPA1^20^. Wild-type and mutant *L. monocytogenes* EGD-e strains were grown as confluent layers on BHI agar plates at 37°C and 42°C and halo diameters around fosfomycin-containing filter discs were measured and corrected for the disc diameter. The experiment was performed in triplicate, and average values and standard deviations are shown. Asterisks indicate statistically significant differences (*P*<0.01).

*Lm*PBPA1^Arg8^ and *Bs*PBP1^Arg8^ are equivalent in their interactions with GpsB. Of the other GpsB-binding determinants of *Bs*PBP1, *Lm*PBPA1 lacks an analogous *Bs*PBP1^Arg11^. The sequential equivalent is *Lm*PBPA1^Tyr11^, but this residue is completely disordered, and its mutation to alanine reduced the affinity for *Bs*GpsB_1–68_^Lys32Glu^ by only 2-fold (**Supplementary Figure 2B**). The importance of α-helix formation in *Lm*PBPA1_1–21_ for GpsB binding was confirmed by CD of *Lm*PBPA1_1–21_^Gln10Pro^ (**Supplementary Figure 2C**) and a concomitant >7-fold reduction in binding affinity (**Supplementary Figure 2B, Supplementary Table 1**). The effects of mutations to the crucial *Lm*GpsB-interacting residues in *Lm*PBPA1 were also probed *in vivo* using fosfomycin sensitivity as a reporter, since *L. monocytogenes* Δ*gpsB* mutants are more susceptible to this antibiotic at 37°C^20^. Effects on fosfomycin sensitivity were apparent in mutants carrying the *pbpA1^Arg8AlaArg12Ala^* and *pbpA1^Gln10Pro^* alleles but only when PBPA2, the PBPA1 paralogue, was also absent (**Figure 2C**). Synthetic lethality with *pbpA2* and a growth defect at 42°C is characteristic of the *L. monocytogenes* null *gpsB* mutant^19^, suggesting that the observed effects partially phenocopy Δ*gpsB*. However, no *pbpA1* mutation completely phenocopied the Δ*gpsB* mutant (**Supplementary Figure 2D**). Taken together, our data highlight the importance of a conserved arginine in class A PG synthases for interacting with GpsB in two species. Furthermore, since *pbpA1* does not phenocopy *gpsB* in *L. monocytogenes*, and *gpsB* deletion on its own in *B. subtilis* has no clear phenotype, GpsB must have additional functions in both bacteria.

### Extending the GpsB interactome in *B. subtilis* and *L. monocytogenes*

The data presented above describe features critical for interactions involving *Bs*GpsB, which include a helical SRxxR(R/K) motif in close proximity to the membrane. To identify hitherto unidentified *Bs*GpsB-interacting proteins, the *B. subtilis* proteome was queried with the SRxxR(R/K) motif. Two previously uncharacterised ORFs, *Bs*YpbE and *Bs*YrrS, conform to all the features described above. *Bs*YpbE is a membrane protein with a 59-residue cytoplasmic domain that encodes a SRVERR motif. The extracellular region, residues 79–240, contains a LysM (lysin motif) domain between residues 189–235; LysM domains are ~40-residue, degenerate PG- and chitin-binding modules widespread in bacteria and eukaryotes. *yrrS* is found in a bicistronic operon widely conserved in the *Bacillaceae* with the gene (*yrrR*) encoding a class B PBP, PBP4b^33^, suggesting these genes have a linked function in cell wall homeostasis^34^. *Bs*YrrS comprises an 18-residue cytoplasmic domain with two potential, overlapping *Bs*GpsB-binding motifs SRYENR and NRDKRR and an extracellular domain that belongs to the widespread and currently uncharacterised DUF1510 family.

LysM domains are frequently found as tandem repeats within bacterial proteins^35^ and the individual domains can act co-operatively to bind PG^36,37^. *Bs*YpbE contains one LysM domain hence oligomerization of *Bs*YpbE may enhance PG binding, with the oligomerisation of the extracellular LysM domain of *Bs*YpbE controlled by cytoplasmic, hexameric *Bs*GpsB^26^, the essential form of the protein *in vivo*^19^. In the absence of purified, full-length *Bs*YpbE to test this hypothesis directly, monomeric and dimeric forms of *Bs*YpbE_130–240_, which encompasses the sole extracellular LysM domain, were generated instead. Dimeric *Bs*YpbE_130–240_ was prepared by mutation of Ser132 to cysteine, enabling disulphide-linked *Bs*YpbE_130–240_^Ser132Cys^ dimers to be purified. In pulldown assays the binding of *Bs*YpbE_130–240_^Ser132Cys^ dimers to PG was enhanced considerably relative to the monomeric, cysteine-free version of *Bs*YpbE_130–240_ (**Supplementary Figure 3A**) and, therefore, the binding of YpbE to PG is stimulated by its multimerisation, presumably driven in *B. subtilis* by hexameric GpsB.

The interaction of *Bs*GpsB_1–68_ with *Bs*YrrS and *Bs*YpbE was assessed by FP and BACTH. *Bs*GpsB_1–68_ bound to *Bs*YpbE_1–21_ and *Bs*YrrS_1–18_ with *K*_d_ values of 13 μM (**Figure 3A**) and 430 μM (**Figure 3B**), respectively. The specificity of these interactions was consistent with the impact of *Bs*GpsB^Asp31Ala^ and *Bs*GpsB^Tyr25Phe^ mutations, each of which reduced the affinities for *Bs*YrrS_1–18_ and *Bs*YpbE_1–21_ by 7- and ~40-fold, respectively (**Figure 3A,3B**), and in-line with the roles of *Bs*GpsB^Asp31^ and *Bs*GpsB^Tyr25^ in defining the *Bs*PBP1 binding site. Interactions of *Bs*GpsB with *Bs*YrrS and *Bs*YpbE were also detected by BACTH, with the interactions mapping to *Bs*GpsB_1–65_ in both cases (**Figure 3C**). GpsB is only conditionally essential in *B. subtilis*^11,12^, and perhaps it is not surprising that no obvious cell growth or division phenotypes could be identified by the deletion of the genes encoding of *Bs*YrrS, *Bs*PBP4b or *Bs*YpbE (data not shown). BACTH was used to confirm that *Bs*YrrS interacted with *Bs*PBP4b, *Bs*PBP1 and *Bs*RodZ; the latter two proteins have established roles in cell division, growth and morphogenesis^38,39^. The *Bs*PBP1:*Bs*YrrS_Δ13–16_ interaction was quantified by SPR, where *Bs*YrrS_Δ13–16_ was used to reduce non-specific binding to the *Bs*PBP1-immobilised SPR chip, and *Bs*YrrS_Δ13–16_ bound to *Bs*PBP1 with a *K*_d_ of 20 nM (**Supplementary Figure 3B**). Therefore, these gene products are capable of forming a network of interactions (**Figure 3D**) that may be nucleated by the formation of a *Bs*PBP1:*Bs*YrrS complex given the affinity of this particular interaction.

**Figure 3.**
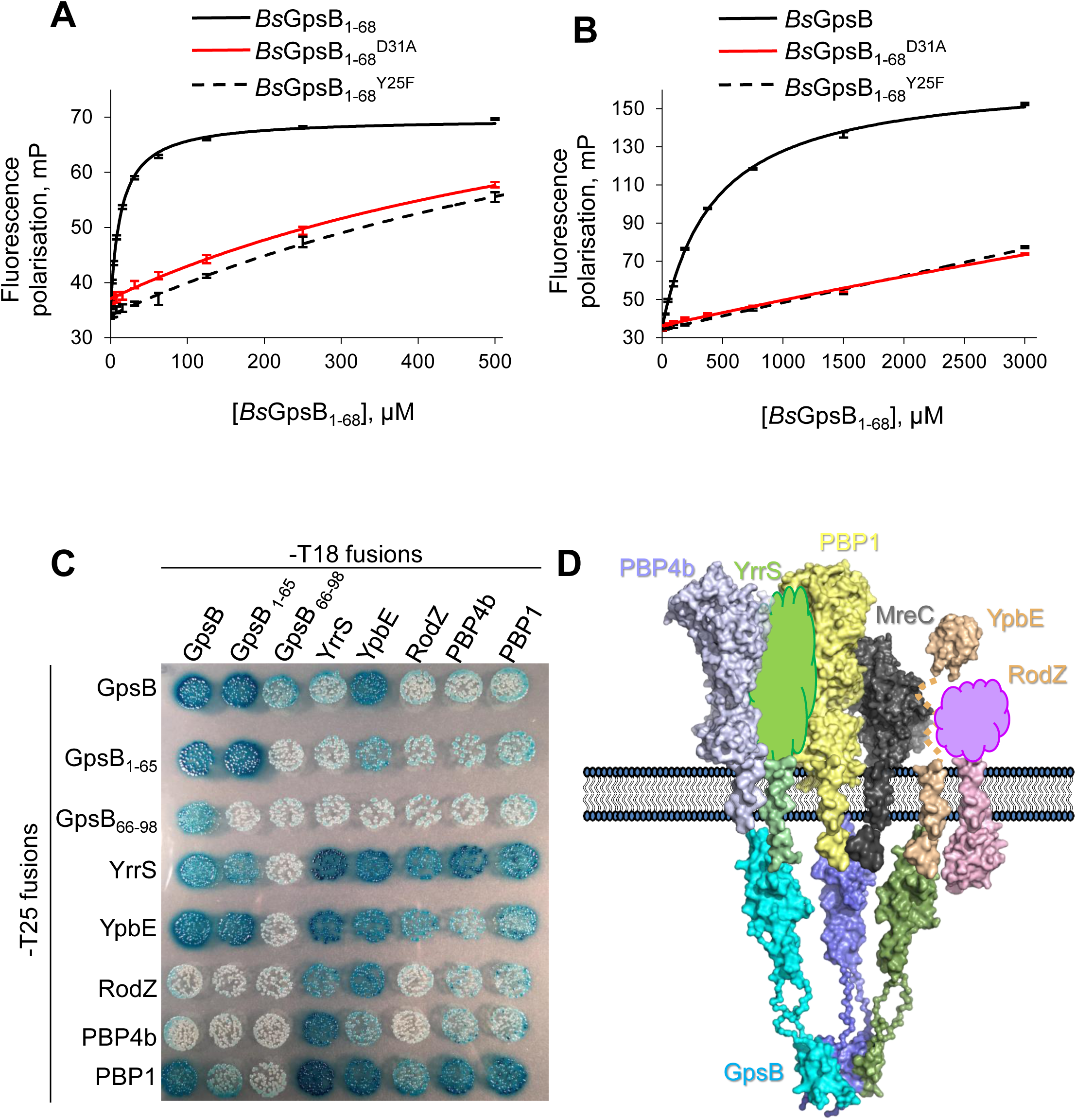
The conserved SRxxR(R/K) motif identifies *Bs*YpbE and *Bs*YrrS as new *Bs*GpsB binding partners. *Bs*YpbE_1–18_ and *Bs*YrrS_1–21_ bind to *Bs*GpsB_1–68_ at the same site as *Bs*PBP1. Fluorescence polarisation of the binding of *Bs*GpsB_1–68_ to fluorescein-labelled *Bs*YpbE_1–18_ (**A**) and fluorescein-labelled *Bs*YrrS_1–21_ (**B**). The interaction of wildtype proteins is depicted by the black curve, whereas the red curve and dashed black line correspond to the same experiment conducted with *Bs*GpsB_1–68_^Asp31Ala^ and *Bs*GpsB_1–68_^Tyr25Phe^ mutants, respectively. (**C**) BACTH reveals a new *Bs*GpsB interaction network involving a set of proteins that encode the conserved SRxxR(R/K) motif. The panel shows pairwise combinations of the proteins expressed as N-terminal fusions to both halves of the adenyl cyclase protein in the BACTH host strain. Their presence in complexes containing *Bs*RodZ, *Bs*PBP4b and *Bs*PBP1 imply roles for *Bs*YrrS and *B*sYpbE in the synthesis of the sidewall during cell growth. (**D**) A model to recapitulate the interactions between *Bs*GpsB *Bs*PBP1, *Bs*PBP4b, *Bs*YrrS, *Bs*YpbE, *Bs*RodZ and *Bs*MreC. GpsB-MreC-PG synthase interactions are common to all three studied species. The individual proteins are coloured separately and each *Bs*GpsB dimer is also coloured independently. Where structural models do not exist, the closest homologue in the PDB has been used instead, or an amorphous blob for where there is no structural information. 18-amino acid model helices represent each TM helix; the predicted N-terminal region of *Bs*PBP4b and *Bs*MreC is only six amino acids and is thus not shown.

Homologues of YpbE do not exist in *L. monocytogenes* and the YrrS homologue (Lmo1495) does not contain a signature *Bs*GpsB-binding motif and neither protein is found in *S. pneumoniae*. No strong potential GpsB-interacting candidates were identified when the *L. monocytogenes* proteome was searched with either TRxxYR or SRxxR(R/K) as the query. BACTH was thus used to uncover additional potential *Lm*GpsB functions in *L. monocytogenes* using a bank of known components from the listerial elongation and division machineries. There is no consensus motif shared by these proteins, though all have at least one arginine present in their cytoplasmic regions that is conserved in their respective orthologues. Two classes of hits were identified in the BACTH screen; class I hits (*Lm*PBPA1, *Lm*MreC and *Lm*SepF, and *Lm*GpsB self-interactions) turned blue after one day of incubation (**Supplementary Figure 3C**). Class II hits turned blue after 2 days incubation at 30°C, including *Lm*ZapA, *Lm*EzrA, *Lm*DivIB, *Lm*DivIC, *Lm*MreC, *Lm*MreBH and the other HMW *Lm*PBPs (**Supplementary Figure 3C**), All of these interactions, except for the GpsB self-interactions, required the *Lm*GpsB N-terminal domain (**Supplementary Figure 3C**). In good agreement with the absence of a TRxxYR motif in *Lm*MreC, *Lm*SepF and *Lm*ZapA, interactions with these proteins did not require key residues in the known PBP-binding groove in *Lm*GpsB (**Supplementary Figure 3D**) and reciprocal tests validated the *Lm*GpsB class I interactions (data not shown). It would thus seem that *Lm*PBPA1 represents the only GpsB binding partner that employs the TRxxYR motif in *L. monocytogenes*.

### A different mode of interaction is used by *S. pneumoniae* GpsB to bind to PBP2a

*S. pneumoniae*, more distantly related to either *Bacillus* or *Listeria* and GpsB, is an ovoid-shaped Gram-positive coccus in which GpsB is essential^22–24^. *Sp*GpsB co-immunoprecipitates with *Sp*PBP2a (one of three pneumococcal class A PBPs), *Sp*MreC and other proteins, suggesting they interact at some point in the pneumococcal cell cycle^24^. Synthetic lethality studies in pneumococcal Δ*gpsB* suppressor mutants revealed that *pbp1a*, and not *pbp2a*, became essential in the absence of *gpsB* indicating that *Sp*PBP2a is the class A PBP regulated by *Sp*GpsB in *S. pneumoniae*^24^. We found that the cytoplasmic mini-domain of *Sp*PBP2a and many of its orthologues contain the consensus sequence (S/R)RS(R/G)(K/S)xR (**Supplementary Figure 4A**) that resembles the *Bacillaceae* PBP1 SRxxR(R/K) motif (**Supplementary Figure 1D**). A peptide encompassing this region, *Sp*PBP2a_23–45_, was found by FP to bind to *Sp*GpsB_1–63_ with a *K*_d_ of 80 μM whereas *Sp*GpsB_1–63_^Asp33Ala^ (equivalent to *Bs*GpsB^Asp35Ala^) had a ~40-fold reduced affinity for *Sp*PBP2a_23–45_ (**Figure 4A, Supplementary Table 1**). The crystal structure of *Sp*GpsB_4–63_ was solved in the presence of *Sp*PBP2a_27–40_; in this instance, each subunit of the *Sp*GpsB dimer is peptide-bound (**Figure 4B**). Peptide binding principally involves two arginines but each *Sp*GpsB subunit recognizes the peptide differently. In *Sp*GpsB_4–63_ molecule 1, *Sp*PBP2a_27–40_ recognition centres on Arg31 and Arg36 (**Figure 4C**), whereas molecule 2 involves Arg33 and Arg36 (**Figure 4D**). The arginine pairs occupy the same positions as *Bs*PBP1^Arg11^ and *Bs*PBP1^Arg8^; Arg36 is equivalent to the former whereas Arg31 or Arg33 are equivalent to the latter.

**Figure 4.**
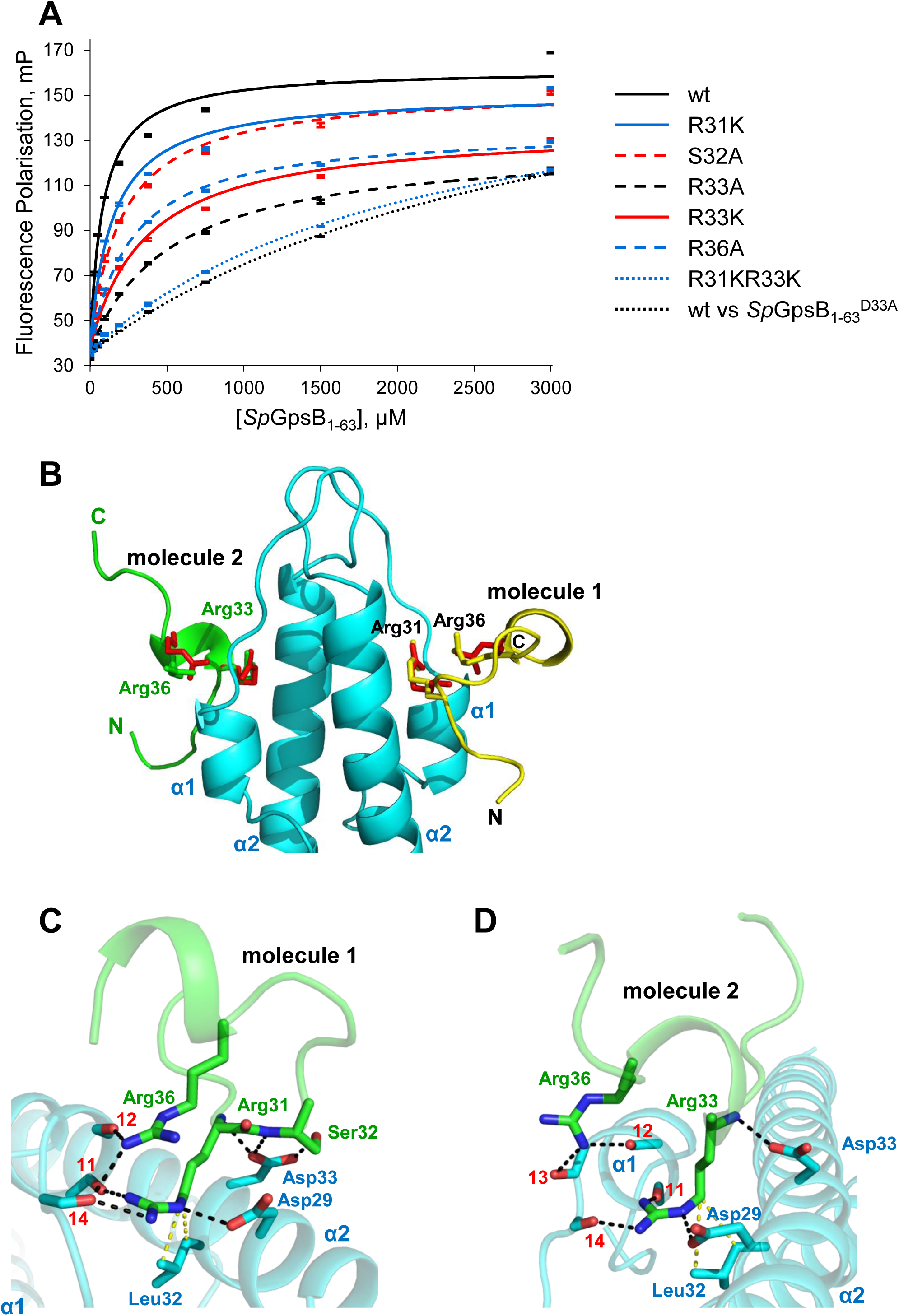
The *Sp*PBP2a minidomain is not α-helical but still interacts with *Sp*GpsB through conserved arginines. (**A**) Arginine residues of *Sp*PBP2a play a key role in binding to *Sp*GpsB. Unless otherwise indicated, the fluorescence polarisation binding curves represent the interaction of TAMRA-labelled *Sp*PBP2a_23–45_ peptides with wildtype *Sp*GpsB_1–63_. The relevant dissociation constants are listed in **Supplementary Table 1**. (**B**) The structure of the *Sp*GpsB_4–63_:*Sp*PBP2a_27–40_ complex reveals the critical role of *Sp*PBP2a arginines for the interaction with *Sp*GpsB. In this cartoon, *Sp*GpsB_4–63_ is coloured cyan, and the *Sp*PBP2a_27–40_ peptide is coloured yellow (molecule 1) and green (molecule 2). The sidechains of Arg8 and Arg11 from the *Bs*GpsB_5–64_:*Bs*PBP1_1–17_ complex are shown as red sticks after a global superimposition of equivalent GpsB atoms. In molecule 1, *Sp*PBP2a^Arg31^ and *Sp*PBP2a^Arg36^ superimpose with *Bs*GpsB_5–64_^Arg8^ and *Bs*GpsB_5–64_^Arg11^ whereas molecule 2 accommodates *Sp*PBP2a^Arg33^ and *Sp*PBP2a^Arg36^. The carbonyl oxygens of *Sp*GpsB^Ile11^, *Sp*GpsB^Phe12^, *Sp*GpsB^Glu13^ and *Sp*GpsB^Gln14^ are denoted by respective red numerals. Close-up view of the interactions of *Sp*PBP2a from molecule 1 (**C**) and 2 (**D**) with *Sp*GpsB_4–63_. Key interfacial sidechains and backbone atoms are represented in stick format; *Sp*GpsB_4–63_ is coloured cyan and *Sp*PBP2A_27–40_ is coloured green. The van der Waals’ interactions between *Sp*GpsB^Leu32^ and *Sp*PBP1^Arg31^ (molecule 1) and *Sp*PBP1^Arg33^ (molecule 2) are in yellow.

The *Sp*GpsB:*Sp*PBP2a interaction was confirmed by BACTH (**Supplementary Figure 4B)**. The *Sp*GpsB:*Sp*PBP2a interaction was lost completely with *Sp*GpsB^Tyr23Ala^, *Sp*GpsB^Val28Ala^, *Sp*GpsB^Asp29Ala^, *Sp*GpsB^Leu32Ala^ and *Sp*GpsB^Asp33Ala^ mutated proteins and reduced with *Sp*GpsB^Ile36Ala^ (**Figure 5A**). All the *Sp*GpsB variants retained the ability to interact with themselves and with wildtype *Sp*GpsB (**Figure 5A, Supplementary Figure 4C**) indicating that these proteins were functional. Moreover, all the *Sp*GpsB variants, except *Sp*GpsB^Asp29Ala^, retained some ability to interact with *Sp*MreC, which was also confirmed to interact with *Sp*GpsB by BACTH (**Figure 5A, Supplementary Figure 4B**). These results indicate that the interface between GpsB and class A PG synthases is conserved in the three models studied here, and the GpsB:MreC interface in *L. monocytogenes* (**Supplementary Figure 3D**) overlaps with that in *S. pneumoniae* (**Figure 5A**).

**Figure 5.**
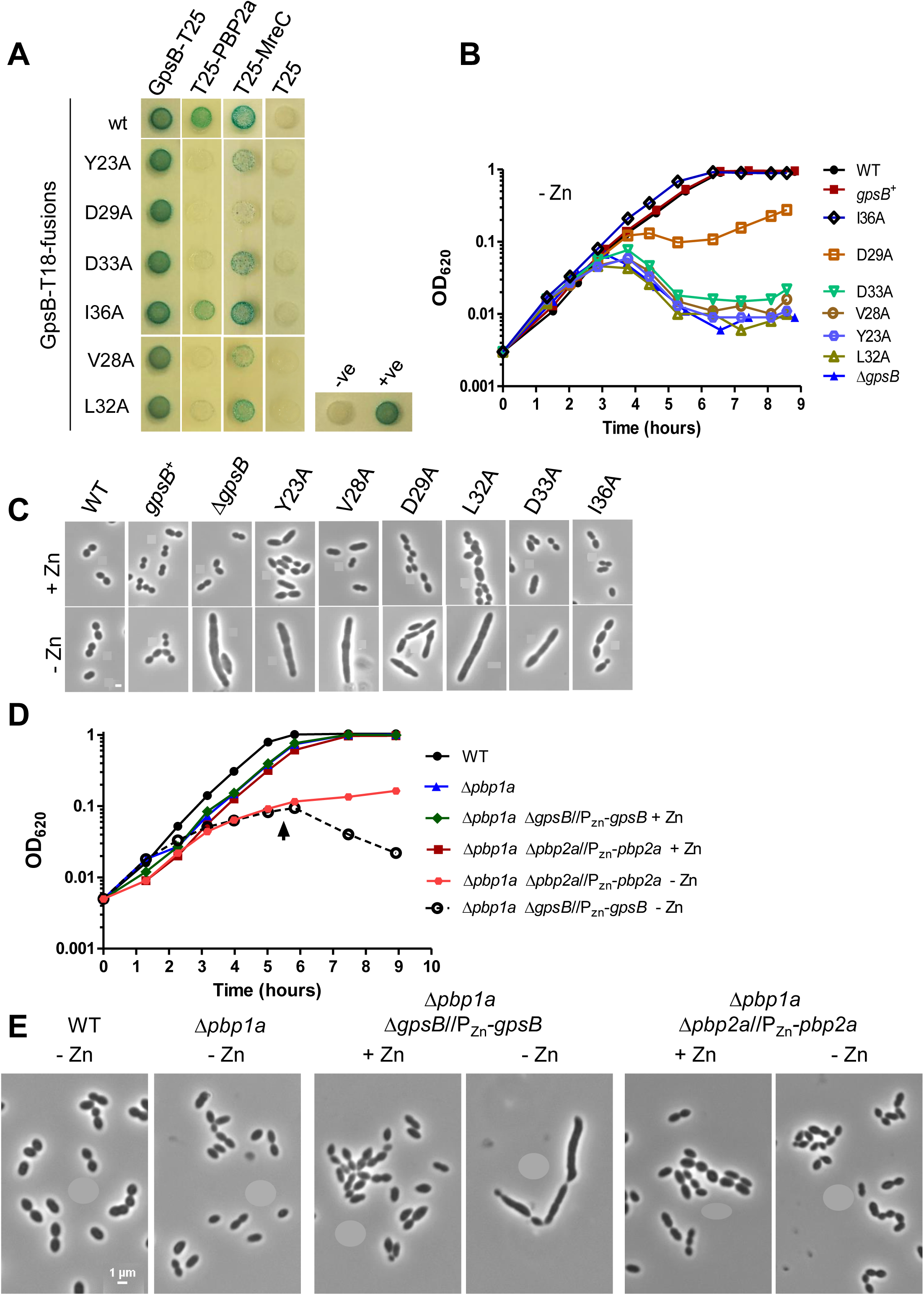
*Sp*PBP2a does not phenocopy *Sp*GpsB. (**A**) Mutations in *Sp*GpsB differentially affect interactions with *Sp*PBP2a and *Sp*MreC. BACTH analysis of the interactions of *Sp*GpsB-T18 variants with wildtype *Sp*GpsB, *Sp*PBP2a and *Sp*MreC. pKT25/pUT18C and pKT25-zip/pUT18C-zip plasmid pairs were used as negative (-ve) and positive (+ve) controls, respectively. The agar plates were photographed after 40 h of incubation at 30°C. *Sp*GpsB variants that have lost *Sp*PBP2a binding have a *gpsB* null growth and morphology phenotype. Representative growth curve of *S. pneumoniae* strains with ectopic expression of *gpsB*^+^ under a Zn^2+^-dependent promoter. GpsB variants of Y23A, V28A, L32A and D33A showed a *gpsB* null growth phenotype (**B**) and elongated cell morphology (**C**) on *gpsB* depletion. The D29A variant showed an intermediate growth phenotype, which was also obtained with an independent isogenic isolate and with a *gpsB*^D29A^-FLAG labelled strain. The I36A strain has a reduced elongation phenotype. All phase-contrast micrographs are at the same magnification (scale bar = 1 μm). *Sp*PBP2a depletion does not phenocopy *Sp*GpsB. Representative growth curves (**D**) and phase-contrast micrographs (**E**) of parent IU1824 (WT, D39 Δ*cps rpsL1*), IU13444 (Δ*pbp1a*), IU14381 (*Δpbp2a//*Δ*bgaA*::P_Zn_-*pbp2a*^+^ Δ*pbp1a*) and IU14383 (*ΔgpsB//*Δ*bgaA*::P_Zn_-*gpsB*^+^ Δ*pbp1a*). Similar to the depletion of *Sp*GpsB in *S. pneumoniae pbp1a*^+^ strains (see panel **B**), depletion of *Sp*GpsB in IU14383 leads to extremely elongated cells, a growth cessation and lysis phenotype. By contrast, depletion of *Sp*PBP2a in the Δ*pbp1a* background (right hand panels) leads to small but mostly ovococcal cells that do not lyse during the time course examined. All phase-contrast micrographs were taken at OD_620_ ≈ 0.15 or at the time point marked by arrows in (A) for IU14381 and IU14383 under zinc depletion and are at the same magnification (scale bar = 1 μm).

Despite differences in the secondary structures of the two independent *Sp*PBP2a peptides bound to the *Sp*GpsB_4–63_ dimer (**Figure 4B**), the two arginines form a similar network of interactions with *Sp*GpsB as described above (**Figure 1C,2A)** with additional sidechain contacts in molecule 1 between *Sp*PBP2a^Ser32^ and *Sp*GpsB^Asp33^ (**Figure 4C**), and *Sp*PBP2a^Arg31^ and *Sp*GpsB^Tyr23^. The importance of *Sp*PBP2a^Arg31^ and *Sp*PBP2a^Arg33^ is further supported by their sequence conservation (**Supplementary Figure 4A**) and FP (**Figure 4A, Supplementary Table 1**). Although *Sp*PBP2a^Arg31Lys^ had only a 2-fold reduced affinity, which probably reflects the ability of Arg33 to compensate for the loss of Arg31, the binding affinity of *Sp*PBP2a^Arg31LysArg33Lys^ was reduced >25-fold relative to wild-type. The importance of the *Sp*GpsB residues involved in the interactions with *Sp*PBP2a is also consistent with the phenotype *in vivo* because of the severe growth (**Figure 5B**) and morphological defects (**Figure 5C**) of *S. pneumoniae* strains harbouring the *Sp*GpsB^Tyr23Ala^, *Sp*GpsB^Val28Ala^, *Sp*GpsB^Asp29Ala^, *Sp*GpsB^Leu32Ala^ and *Sp*GpsB^Asp33Ala^ alleles even though the mutated proteins were capable of self-interactions (**Supplementary Figure 4C**) and were expressed at wildtype levels (**Supplementary Figure 4D**). However, no obvious phenotype was observed in *S. pneumoniae* strains carrying the corresponding *Sp*PBP2a^Arg31Ala^, *Sp*PBP2a^Arg31Lys^, *Sp*PBP2a^Arg33Ala^ or *Sp*PBP2a^Arg31AlaSer32AlaArg36Ala^ alleles, even when *pbp1a* was deleted to decouple the effects of mutations in *Sp*PBP2a from *Sp*PBP1a activity (**Supplementary Table 2**, data not shown). Nevertheless, *Sp*PBP2a mutants in which amino acids 32–37 or 27–38 or 26–45 were deleted in a Δ*pbp1a* background showed progressively reduced growth rates in the three deletion strains and pronounced morphological defects in the two strains with large deletions (**Supplementary Figure 5A, 5B**), despite wildtype levels of protein expression (**Supplementary Figure 5C)**. BACTH results for the correspondent truncated *Sp*PBP2a variants showed reduced interactions with *Sp*GpsB in comparison to the wildtype but not with *Sp*MreC (**Supplementary Figure 5D, 5E**). Together, these results support a critical role *in vivo* of the (S/R)RS(R/G)(K/S)xR motif between *Sp*PBP2a residues 30 and 36. However, the observation that all three Δ*pbp1a pbp2a* deletion mutants are viable and both growth and morphology phenotypes are different between *S. pneumoniae* Δ*pbp1a* strains depleted for *gpsB* (**Figure 5D**) and Δ*pbp1a* strains depleted for *pbp*2a (**Figure 5E**) implies that *Sp*PBP2a binding is just one function for GpsB in *S. pneumoniae*.

Taken as a whole, our data on three important bacterial systems agree that GpsB is an adaptor protein that connects a major class A PG synthase with other cell wall and cell cycle proteins, and to cell shape determinants such as MreC. The identity and mode of interaction of the GpsB-binding partners varies from species to species and may reflect the different physiologies of each bacterium and their modes of growth and division.

## Discussion

Bacterial cell growth and division necessitates tight co-ordination between the replication and segregation of the chromosome, the fission of the cell membrane and the remodelling of the PG. Consequently proteins and their complexes with major functions on either side of the membrane must co-ordinate their activities. One potential mechanism involves the interactions of major PG synthases with their intracellular regulators. Herein we present the first structures of the cell cycle adaptor, GpsB, in complex with the cytoplasmic mini-domains of PG synthases from three different bacteria, the rod-shaped *B. subtilis* and *L. monocytogenes*, in which *gpsB* is conditionally essential^11,12,19^ and the ovococcal *S. pneumoniae* in which *gpsB* is essential^22–24^. In common with mammalian adaptors GGA^30^ and 14–3-3^40^ proteins, the primary binding surface of GpsB is restricted to a conserved groove between α-helices. The cytoplasmic mini-domains of the three PG synthases in the three organisms have little in common except that each utilises a conserved arginine in their respective sequences to interact with the cognate GpsB. The PG synthase arginine ‘finger’ pokes into a negatively-charged cavity situated between α-helices 1 and 2 of GpsB and is fixed in the same orientation in all structures, just as the phosphoryl group defines the binding orientation of peptides to 14-3-3^40^. The arginine complements the cavity best when the mainchain amide protons of it and its downstream residue are accessible to form hydrogen bonds with *Bs*GpsB^Asp35^, *Lm*GspB^Asp37^ or *Sp*GspB^Asp33^. This scenario can occur when the arginine is either at the start of an α-helix, such as *Bs*PBP1^Arg8^, or at the *i*+1 position in a type I β-turn, such as *Sp*PBP2a^Arg31^, and the bidentate nature of this interaction explains why free L-arginine does not displace pre-bound PBP peptides from GpsB even when present at 100-fold molar excess (data not shown). Similarly, contact to the backbone amide at the *i*+2 position in 14-3-3 ligands is essential for binding^40^. Despite a lack of strong sequence and structural homology in the PG synthase cytoplasmic mini-domains, their binding is dependent upon an identical subset of GpsB residues including *Bs*GpsB^Tyr25^, *Bs*GpsB^Asp31^, *Bs*GpsB^Asp35^ (**Figure 1C**) and their structural equivalents *Lm*GspB^Tyr27^, *Lm*GspB^Asp33^, *Lm*GspB^Asp37^ (**Figure 2A**), and *Sp*GpsB^Tyr23^, *Sp*GpsB^Asp29^ and *Sp*GspB^Asp33^ (**Figure 4C,D**). These amino acids are also conserved in the DivIVA/Wag31/antigen 84 actinobacterial homologues of GpsB, suggesting a role for them in recruiting cell wall synthesis enzymes to the hyphal tip and future branch sites^13,14^, regions that require nascent PG synthesis in filamentous bacteria.

We originally set out to establish the common rules by which GpsB interacts with major PG synthases. Other than the arginine finger mode of GpsB recognition, we discovered how GpsB interacts with at least one class A PBP in each species; that *Lm*GpsB interacts with both the cell shape determinant, MreC41, and a regulator of Z-ring dynamics, EzrA^42^, and confirmed the *Sp*GpsB:MreC interaction by BACTH. These new data complement what was previously known about these interactions in *B. subtilis*^11^ and *S. pneumoniae*^23,24^. How GpsB can interact with such disparate targets remains unknown but *Bs*GpsB^Asp31^, *Lm*GpsB^Asp33^ and *Sp*GpsB^Asp29^ are important for interactions with PBPs and other proteins, including MreC, while *Lm*GpsB^Asp37^ and *Sp*GpsB^Asp33^ only interact with PBPs. There must be at least one other surface that is used by GpsB to form complexes with other proteins in its function as an adaptor.

The GpsB:PBP interaction interface notably requires no more than three sidechains from any PBP to complex with GpsB. Protein:peptide contacts involving less well-conserved exosites that flank a small core, conserved peptide motif can contribute significantly to the affinity of protein:peptide interactions^43,44^, and the contribution of exosites to affinity may explain why point mutations in *Lm*PBPA1 and *Sp*PBP2a have a significant impact using peptide fragments *in vitro*, but have reduced impact *in vivo*. For instance, the *Lm*PBPA1^Arg8Ala^ mutation had negligible effect *in vivo* (**Figure 2C**, **Supplementary Figure 2D**) yet it reduced binding by >15-fold (**Supplementary Table 1**). Similarly *Sp*PBP2a^Arg31LysArg33Lys^ had a >25-fold impact on binding (**Supplementary Table 1**) yet growth or morphology phenotypes were not evident (**Supplementary Table 2**, data not shown) until significant stretches of the cytoplasmic mini-domain were deleted (**Supplementary Figure 5B**).

We also found some differences in GpsB interactions between the species that may be related to GpsB species-specific function. In *B. subtilis*, we discovered a critical motif, SRxxR(R/K), found in close proximity to the membrane that could be used to predict novel GpsB partners. We used this information to identify an interaction network involving cell envelope binding and modifying proteins that most likely is underpinned by the GpsB hexamer. An RSxxxR motif was identified in class A PBPs from most streptococci and sequence features that could dictate GpsB-binding can be found in class A PBPs in other Gram-positive organisms such as the lactococci*, Leuconostocaceae* and enterococci, including the ESKAPE pathogen *E. faecium* (**Supplementary Figure 6**). However, sequence-based searches alone will not identify complete GpsB interactomes because the local structure of the sequence and its proximity to the membrane are also key parameters of GpsB binding. *Sp*PBP2a contains a partially conserved sequence RSxxxR (**Supplementary Figure 4A**) that resembles the *Bs*PBP1 signature motif and the *Sp*PBP2a mini-domain interacts with each subunit of the *Sp*GpsB dimer in a different way (**Figure 4B-D**). Finally, we have observed here that mutation or deletion of the cytoplasmic mini-domain of *Sp*PBP2a does not phenocopy deletion or depletion of *gpsB* (**Figure 5C, 5E**). Similarly, a Δ*pbpA1* strain did not phenocopy *gpsB* deletion in *L. monocytogenes*^19^. Taken together these results imply that there must be at least one other critical GpsB interaction partner, beyond respective class A PG synthases, that dictates its conditional essentiality in *L. monocytogenes* and *B. subtilis*, and essentiality under normal growth conditions in *S. pneumoniae*.

In every Firmicute (and Actinobacterium) tested thus far, GpsB (or the homologous DivIVA/Wag31/antigen 84) acts as an adaptor to co-ordinate PG synthase activity with other processes depending on the physiology of the cell. GpsB hexamerisation can thus bridge the interaction of multiple binding partners, a function GpsB shares with 14-3-3 proteins that can form ternary complexes with BCR and Raf-1 by 14-3-3 dimerisation^45^. In bacilli, *Bs*GpsB plays a role in shuttling between the side wall during elongation and the septum during division^11^ and, given that *Bs*PBP4b is regulated by σ factors E^33^ and F^46^, complexes of *Bs*PBP4b and *Bs*YrrS, bridged by *Bs*GpsB (**Figure 3C,D**), presumably play a role in the asymmetric cell division characteristic of endospore-forming bacilli. In listeria, which is closely related to bacilli and shares with them a rod-like morphology, GpsB appears to connect several PBPs with proteins with known roles in cytokinesis, including Z-ring polymerisation modulators (ZapA, EzrA, SepF), late division proteins (DivIB, DivIC) and the elongasome (MreC, MreBH) (**Supplementary Figure 3C,D**), all of which except SepF and MreBH have also been tested in *B. subtilis* and found not to interact with *Bs*GpsB^11^.

By contrast, the pneumococci have an ovoid cell shape and lack key components such as the MinCD system for cell division site selection^47^, and MreB-like proteins required for side wall synthesis^48^. Presumably *Sp*GpsB interacts with one or more pneumococcal-specific proteins, the loss of which may be related to the lethal phenotype. Furthermore, *Sp*GpsB affects both StkP autophosphorylation^24,49^ and the StkP-catalysed phosphorylation of *Sp*DivIVA^24,49^, *Sp*MapZ/LocZ^24,50^, *Sp*Jag/EloR/KhpB^51–53^ and *Sp*MacP^54^. It is not yet clear how the complexes formed by these proteins are affected by their phosphorylation, except that *Sp*PBP2a activity is dependent upon phosphorylated *Sp*MacP^54^, at least in the presence of functional *Sp*StkP, or what the impact is of potential cross-talk to two-component signalling systems^55^.

Finally, the different phenotypic outcomes associated with *gpsB* deletion or depletion in the three systems studied herein may reflect the presence of redundant systems in the large genome (4.2 Mbp) of the bacilli, partial redundancy in listeria (2.9 Mbp), and a relative absence of redundancy in the stripped-down genome (2.1 Mbp) of the pneumococci. The relative affinities and cellular concentrations of GpsB partners probably dictate which protein(s) is bound by GpsB at any point in a cell cycle-dependent manner; simultaneous interactions with multiple target proteins is likely to lead to an increase in avidity of GpsB as more commony found in antibody:antigen interactions. However, the intricate networks involving GpsB will only be uncovered by validating the full GpsB interactome.

## Materials and Methods

Full details of all the experimental procedures are presented in the **Supplementary Information**.

### Bacterial strains and growth conditions

**Supplementary Table 3** lists all bacterial strains used in this study that were grown in brain heart infusion (BHI) or LB media at 37°C (or 42°C, where indicated), supplemented with antibiotics if required. Standard *Escherichia coli* strains were used as cloning^56^ and recombinant protein production hosts.

### General methods, manipulation of DNA and oligonucleotide primers

Bacterial transformation and isolation of plasmid DNA, enzymatic DNA modification and Quikchange mutagenesis was performed according to standard protocols^56–58^ using the plasmids in **Supplementary Table 4** and the oligonucleotide primers in **Supplementary Tables 5–7**. All constructs were verified by Sanger DNA sequencing.

### Bacterial two-hybrid experiments (BACTH)

The BACTH system^59^ was used to screen GpsB proteins against potential binding partners. The genes of interest were amplified by PCR and cloned by restriction into the appropriate BACTH vectors to obtain the corresponding N- and C-terminal hybrid fusions. Domain deletions and single point mutations were introduced by Quikchange or by subcloning the specific mutated alleles amplified from their respective DNA template. Plasmids encoding the respective genes fused to the N- or C-termini of the T18- or the T25-fragment of the *Bordetella pertussis* adenylate cyclase were co-transformed into *E. coli* BTH101. Co-transformants were selected on nutrient or LB agar plates containing ampicillin (100 μg/mL), kanamycin (50 μg/mL), X-Gal (40 μg/mL) and IPTG (0.1 - 0.5 mM). Photographs were taken after at least 24 h of incubation at 30°C.

### Construction of *L. monocytogenes* and *S. pneumoniae* mutant strains

Full details of the strain construction can be found in the **Supplementary Information**.

### *L. monocytogenes* fosfomycin susceptibility assays

Fosfomycin susceptibility of *L. monocytogenes* was measured, and corrected for the disc diameter, after overnight incubation using fosfomycin-impregnated filter discs.

### Recombinant protein and peptide production

Recombinant expression constructs for *Bs*GpsB_5–64_, *Sp*GpsB_1–63_ and *Sp*GpsB_4–63_ were prepared along similar lines as those for *Bs*GpsB_1–68_ and *Lm*GpsB_1–73_, as described previously^19^. Maltose binding protein (MBP) fusions of *Lm*PBPA1_1–20_ and *Sp*PBP2a_23–45_ were cloned into pMAT11, a modified version of pHAT4^60^, as described previously for MBP- *Bs*PBP1_1–32_^19^. The ORF encoding *Bs*YpbE_80–240_ was cloned into pET28a before the DNA encoding residues 80–129 was deleted by PCR amplification of the entire plasmid to yield *Bs*YpbE_130–240_. The construct for expressing *Bs*YrrS_Δ13–16_ was prepared by cloning *yrrS* into pET28a and the DNA coding for residues 13–16 was deleted by Quikchange.

*Bs*GpsB_1–68_, *Sp*GpsB_1–63_ and *Lm*GpsB_1–73_ were purified as described previously^19^. *Sp*GpsB_1–63_ was further purified by ammonium sulphate precipitation after thrombin removal of the His_6_-tag. *Bs*GpsB_5–64_, the *Bs*GpsB_5–64_^K32E^ mutant and *Sp*GpsB_4–63_ were expressed and purified by a similar protocol except that TEV protease, rather than thrombin, was used to remove the N-terminal His_6_-tag by overnight cleavage. YpbE_130–240_ and its Ser132Cys variant were expressed in *E. coli* BL21(DE3) and purified by ion exchange, ammonium sulphate precipitation and size exclusion. YrrSΔ_13–16_ was expressed in *E. coli* BL21(DE3) and His^6^-tagged YrrSΔ_13–16_ was purified from the membrane fraction by Ni^+^-NTA affinity and size exclusion chromatography. The PBP peptides, generated as MBP-fusion proteins, were expressed, purified, fluorescently-labelled and separated from the MBP fusion partner as described previously^19^.

All recombinant proteins or peptides were concentrated and flash frozen in small aliquots in liquid nitrogen and stored at -80°C.

The *Bs*PBP1_1–17_, *Lm*PBPA1_1–15_ and SpPBP2a_27–40_ peptides were synthesized chemically (Protein and Peptide Research Ltd, UK and Severn Biotech, UK).

### Crystallization and structure determination

Co-crystallization of *Bs*GpsB_5–64_:*Bs*PBP1_1–17_, *Bs*GpsB_5–64_^Lys32Glu^:*Lm*PBPA1_1–15_ and *Sp*GpsB_4–63_:*Sp*PBP2a_27–40_ followed the same procedure. Equal volumes of GpsB protein and PBP peptide were mixed at final concentrations of 20 and 25 mg/mL, respectively, corresponding to a 1:5 molar ratio. The protein:peptide complexes were crystallised at room temperature using commercial crystallization screens and a Mosquito (TTP Labtech) liquid handling robot. All crystals were mounted in rayon loops and frozen directly in liquid nitrogen.

All diffraction data were collected at beamlines I24 and I04 at the Diamond synchrotron light source except for unbound *Sp*GpsB_4–63_, which were collected in house using a gallium METALJET™ X-ray source (Bruker AXS GmbH). Diffraction images for *Bs*GpsB_5–64_:*Bs*PBP1_1–17_ and *Sp*GpsB_4–63_:*Sp*PBP2a_27–40_ were indexed and integrated with XDS^61^ and scaled with AIMLESS^62^; for *Bs*GpsB_5–64_^Lys32Glu^:*Lm*PBPA1_1–17_ indexing and integration took place with DIALS^63^, scaling in XDS^61^ and merging with AIMLESS^62^. For unbound *Sp*GpsB_4–63_ the images were indexed, integrated, scaled and merged with Proteum 3 (Bruker AXS GmbH). All structures were solved by molecular replacement in PHASER^64^ and rebuilt in COOT^65^, interspersed with rounds of refinement in REFMAC^66^ and PHENIX.REFINE^67^. Statistics for data collection and the final refined models can be found in **Supplementary Table 8**.

### FP assays

FP experiments were undertaken as described previously^19^, in a buffer of 10 mM Tris.HCl (pH 8.0), 250 mM NaCl, 0.1% reduced Triton X-100. Where TAMRA-labelled peptides were used in FP assays the excitation wavelength was 540 nm and fluorescence emission was recorded above 590 nm.

### SPR

All SPR experiments used a running buffer of 10 mM Tris.HCl (pH 8.0), 250 mM NaCl, 0.1% reduced Triton X-100. *Bs*PBP1 and *Bs*PBP1_17–914_ were immobilized on the surface of a CM5 chip (GE Healthcare) as described previously^19^. For the PBP1/YrrS_Δ13–16_ titration 800 RUs of *Bs*PBP1 were immobilized on the chip surface; for the *Bs*PBP1/*Bs*GpsB titration 1200 RUs of *Bs*PBP1 were immobilized.

### CD analysis

CD spectra were recorded on a JASCO J-810 spectropolarimeter with a PTC-4235 Peltier temperature controller using 1 mm path length quartz cuvettes. For full wavelength scans a scan speed of 10 nm/min and a response time of 4 s were used, the final spectra were the average of 4–5 measurements. For thermal melt experiments, a response time of 8 s and scan rate of 1°C/min were used.

### Peptidoglycan pulldown assay

PG pulldown assays were carried out in PBS buffer. *B. subtilis* PG (SigmaAldrich) was prepared as a 10 mg/mL stock. 25 μg of protein were added to 66 μg of PG and incubated for 30 minutes. After two centrifugation and resuspension steps the final pellet was boiled in SDS-PAGE loading buffer before analysis by SDS-PAGE without reducing agents.

## Acknowledgements

This work was supported by grants from the UK BBSRC (BB/M001180/1 to R.J.L.), the German DFG (HA 6830/1-1 and HA 6830/1-2 to S.H.), the American NIH (RO1GM113172 and RO1GM114315 to M.E.W). O.M. is funded by a Contribution to Basic Research (LR7/2007) from the Autonomous Region of Sardinia. F.C. is funded by the European Commission (International Training Network Train2Target, No. 721484) and Z.R. is funded by a UK BBSRC DTP studentship (BB/M011186/1). We thank the Diamond synchrotron light source for access to its beamlines and thank its staff for support during data collection. We are indebted to Waldemar Vollmer for critical reading of the manuscript and for supplying *S. pneumoniae* genomic DNA. We thank Helen Waller for technical assistance with SPR experiments and Simon Thorpe at the University of Sheffield for mass spectroscopy analysis of peptide samples.

